# Rostral Pedunculopontine Nucleus Infusion of M_4_ Positive Allosteric Modulator VU0467154 Augments L-DOPA Effects in Hemiparkinsonian Rats

**DOI:** 10.1101/2024.04.16.589785

**Authors:** Nicole E. Chambers, Annique McLune, Michael Coyle, Ashley Centner, Jordan Sergio, Isabella Delpriore, Kathryn Lanza, Craig W. Lindsley, P. Jeffrey Conn, Christopher Bishop

## Abstract

Standard treatment for Parkinson’s disease (PD) is dopamine replacement therapy with L-DOPA. However, chronic treatment often results in abnormal involuntary movements called L-DOPA-induced dyskinesia (LID). Prior evidence indicates that heightened striatal cholinergic tone may contribute to LID. Restoring cholinergic inhibition by targeting the inhibitory M_4_ muscarinic acetylcholine (ACh) receptor (M_4_) reduces LID in preclinical models. Although intrinsic striatal sources of ACh have been considered for their role in LID, extrinsic sources of ACh such as the pedunculopontine nucleus (PPN) have not been well investigated for their role in LID. Therefore, the current study employed hemiparkinsonian Long-Evans rats with a PPN-targeted cannula ipsilateral to 6-OHDA lesion. Following chronic treatment with L-DOPA, we examined the effect of local unilateral PPN infusion of M_4_ PAM VU0467154 on LID, motor performance, and c-fos expression within the PPN. It was expected that PPN infusion of VU0467154 would reduce LID, reduce L-DOPA’s motor benefit, and globally reduce c-fos expression in the PPN. Contrary to our expectations, PPN infusion of M_4_ PAM did not significantly affect LID severity. Furthermore, the group receiving M_4_ PAM showed slightly elevated motor improvement compared to L-DOPA, and decreased c-fos expression specifically in PPN cholinergic neurons. These results suggest that local PPN ACh dynamics differ from those of the striatum. Specifically, our results suggest that PPN cholinergic neurons may be a promising therapeutic target for augmenting L-DOPA-mediated motor benefit without increasing LID.

## 1. Introduction

Gold standard treatment for Parkinson’s disease (PD) is dopamine (DA) replacement with L-DOPA. However, after chronic L-DOPA treatment most PD patients develop debilitating abnormal involuntary movements called L-DOPA-induced dyskinesia (LID) (Ahlskog & Muenter, 2001). Although causes of LID are multifaceted, heightened striatal cholinergic tone has been linked to an increase in LID severity (Conti et al., 2018; Ding et al., 2010; 2011; Won et al., 2014; among others). One way to normalize cholinergic tone in LID is by potentiating activity of the Gα_i/o_-coupled muscarinic acetylcholine (ACh) M_4_ receptor (M_4_) using an M_4_ positive allosteric modulator (PAM). Indeed, systemic administration of M_4_ PAM VU0467154 has been shown to reduce LID severity in preclinical PD models (Shen et al., 2015). M_4_ are located on both dendrites (Shen et al., 2015) and axons (Moehle et al., 2017) of DA D_1_-bearing medium spiny neurons (MSNs). This location allows M_4_ to oppose DA signaling and decrease activity of the direct pathway which is overactive in LID (Chambers et al., 2023). Targeting M_4_ likely also reduces ACh release, both from intrinsic striatal sources such as cholinergic interneurons, but also from extrinsic sources such as the rostral pedunculopontine nucleus (rPPN; Dautan et al., 2014; 2016). Importantly, cholinergic neurons in the rostral pedunculopontine nucleus project to brain areas critical for motor behavior such as the striatum, substantia nigra pars reticulata, and substantia nigra pars compacta, (Dautan et al., 2014; Moehle et al., 2017; Xiao et al., 2015), whereas caudal PPN cholinergic neurons project to brain areas important for reward learning (Dautan et al., 2014; Xiao et al., 2015). Therefore, even though more cholinergic neurons are expressed in the caudal PPN, the projections from cholinergic neurons in the rostral PPN are most likely to influence motor behavior.

It is important to consider the role of the PPN in LID, because increasing evidence suggests that PPN cholinergic neurons contribute to drug-induced stereotyped movements (Bastide et al., 2014; Chambers et al., 2019; 2021; Inglis et al., 1994; Mathur et al., 1997). It is equally important to consider how the PPN may influence motor benefit derived from L-DOPA. Research by Pienaar and colleagues (2015; Sharma et al., 2020) shows that chemogenetic stimulation of PPN cholinergic neurons improves forepaw akinesia in parkinsonian rodent models. Human clinical literature also suggests that stimulation of PPN neurons may augment the effects of L-DOPA as PD patients receiving chronic PPN deep brain stimulation are able to reduce their dose of L-DOPA (Mazzone et al., 2014). Literature examining the relationship between immediate early gene (IEG) expression and LID also suggests that the PPN may contribute to L-DOPA-mediated effects. The IEG ΔFosB shows increased expression in response to cellular activation. Importantly, increased IEG expression in certain brain areas such as the striatum has been linked to LID, suggesting an upregulation of neural activity as a result of L-DOPA treatment (Bastide et al., 2014; Cenci et al., 2002; McClung et al., 2004; Soghomonian 2006; Sgambato-Faure et al., 2005). Expression of ΔFosB is increased in the PPN of dyskinetic rodents (Ding et al., 2007; Bastide et al., 2014). This suggests that PPN activity may contribute to both LID and L-DOPA’s anti-PD efficacy, however this assertion remains uninvestigated. Therefore, we tested whether local PPN administration of the M_4_ PAM VU0467154 would alter dyskinesia and L-DOPA-mediated motor improvement in hemiparkinsonian rats. Furthermore, using c-fos as an indicator of cellular activation in the PPN, we performed dual immunofluorescent immunohistochemistry to test whether the activity of PPN cholinergic neurons is altered by L-DOPA administration. We hypothesized that PPN infusion of the M_4_ PAM VU0467154 would lessen LID severity and globally decrease PPN c-fos expression, as PPN cholinergic, glutamatergic, and GABAergic neurons all likely contain M_4_ (Langmead et al., 1995; Vilaró et al., 1994; 1995; Ye et al., 2010). We further hypothesized that targeting M_4_ with a PAM would result in a reduction in L-DOPA’s motor efficacy and that PPN cholinergic neuron activity would be elevated by L-DOPA administration, suggesting a role for PPN neurons in modulating motor benefit with L-DOPA.

## 2.0 Experimental Procedures

### 2.1 Subjects

Male and Female Long-Evans rats (N = 28 included in final analyses, N = 54 total; see Figure 1 for numbers in each group) were maintained in a colony room at 22-23° C with a 12h light/dark cycle (lights on at 0700) and ad libitum access to food and water. They were group-housed until surgery, after which they were single-housed in plastic cages (25 cm high, 45 cm deep, 23 cm wide) to prevent damage to the implanted cannula. All experiments were performed in accordance with policies set forth by the Institutional Animal Care and Use Committee at Binghamton University and the current edition of the NIH Guide for Care and Use of Laboratory Animals. Cannula placement was histologically verified in the PPN and DA lesion was verified via tyrosine hydroxylase (TH) immunohistochemistry for each animal included in the analyses.

**Figure 1.**
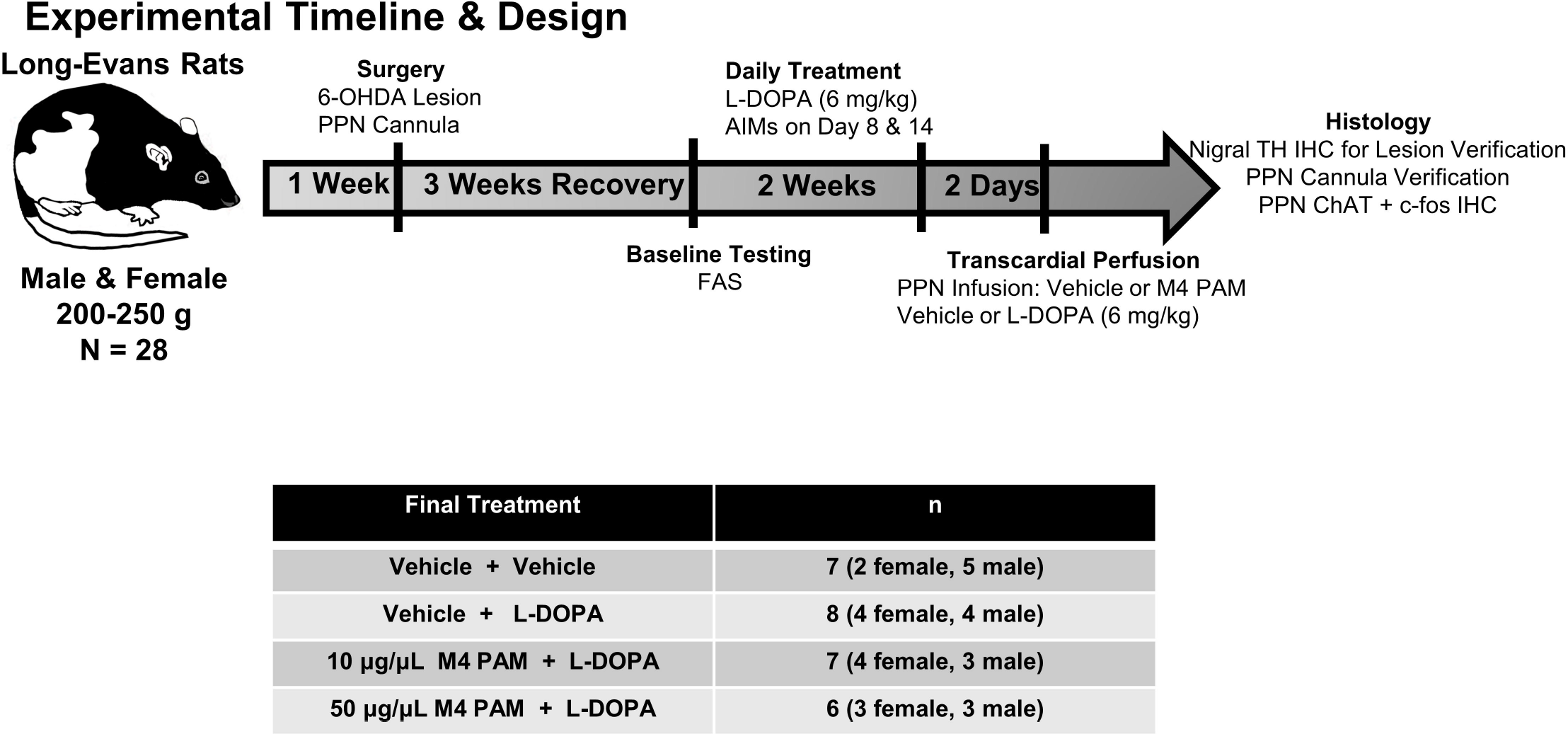
Experimental Design. Male and Female Long-Evans Rats (N = 28; Vehicle + Vehicle = 7; Vehicle + L-DOPA = 8; VU0467154 10 µg/µL + L-DOPA = 7, VU0467154 50 µg/µL = 6) received unilateral medial forebrain bundle lesion with 6-hydroxydopamine (6-OHDA) plus a microinjection cannula implant above the pedunculopontine nucleus (PPN). After 3 weeks of recovery, we measured lesion effects on akinesia using the forepaw adjusting steps test (FAS). After this initial baseline testing, rats received two weeks of daily L-DOPA treatment (6 mg/kg s.c.). During days 8 and 14 of treatment dyskinesia was monitored using the abnormal involuntary movements scale (AIMs). Next, rats were divided into counterbalanced groups with equal levels of motor impairment and dyskinesia. Following 2 days of washout, rats received a PPN-targeted microinfusion of vehicle (100% dimethylsulfoxide), VU0467154 10 µg/µL, or VU0467154 50 µg/µL. Five min after the start of the microinfusion rats were given either vehicle (0.1% ascorbate in 0.9% NaCl) or L-DOPA (6 mg/kg, s.c.). Next, dyskinesia was rated using the validated AIMs scale every 10 min for 90 min. Rats were transcardially perfused after the 90 min rating and brains were harvested for later nigral tyrosine hydroxylase (TH) immunohistochemistry (IHC) to verify successful lesion, as well as co-staining for choline acetyltransferase (ChAT) and c-fos to examine cellular activity in PPN cholinergic neurons. Successful placement of injections in the PPN was verified in all rats. For all PPN histology we did our best to choose slices just outside of the location of the infusion as the slices within the boundary of the microinfusion because slices within this region sustained considerable damage. Rats were retained for these analyses only if they had a DA lesion (< 35% TH percent intact), and if they had successful placement of the infusion in the rPPN.

### 2.2 Procedure

Three weeks after surgery, rats were tested off treatment for baseline motor deficits via forepaw adjusting steps (FAS; See Figure 1 for a detailed experimental timeline). Next, rats received daily L-DOPA treatment (6 mg/kg, s.c.) for 2 weeks to induce stable dyskinesia (Chambers et al., 2019; 2021). The Abnormal Involuntary Movements scale (AIMs) was used to rate dyskinesia. AIMs and FAS were assayed on days 8 and 14. Median AIMs and FAS scores were used to form equally lesioned and dyskinetic counterbalanced treatment groups for the final day of treatment (See Figure 2A for dyskinesia, and 2B for baseline forepaw stepping). After 2 days of drug washout, rats received PPN-targeted infusion of vehicle (100% DMSO), 10, or 50 µg/µL VU0467154 followed 5 min later by L-DOPA (6 mg/kg, s.c.). AIMs ratings occurred every 10 min after L-DOPA administration for 90 min (Figure 2C), and FAS was assayed at the 60 min time point (Figure 2D). After 90 min, rats were taken to the procedure room where they were injected with a lethal dose of pentobarbital and transcardially perfused with ice-cold saline followed by 4% paraformaldehyde (Chambers et al., 2021). Brains were harvested and processed for eventual nigral TH immunohistochemistry (IHC, Figure 3A-C; Table 1), PPN cannula verification (Figure 3D-E), and PPN choline acetyltransferase (ChAT) and c-fos IHC (Figure 4A-B, Table 2).

**Figure 2.**
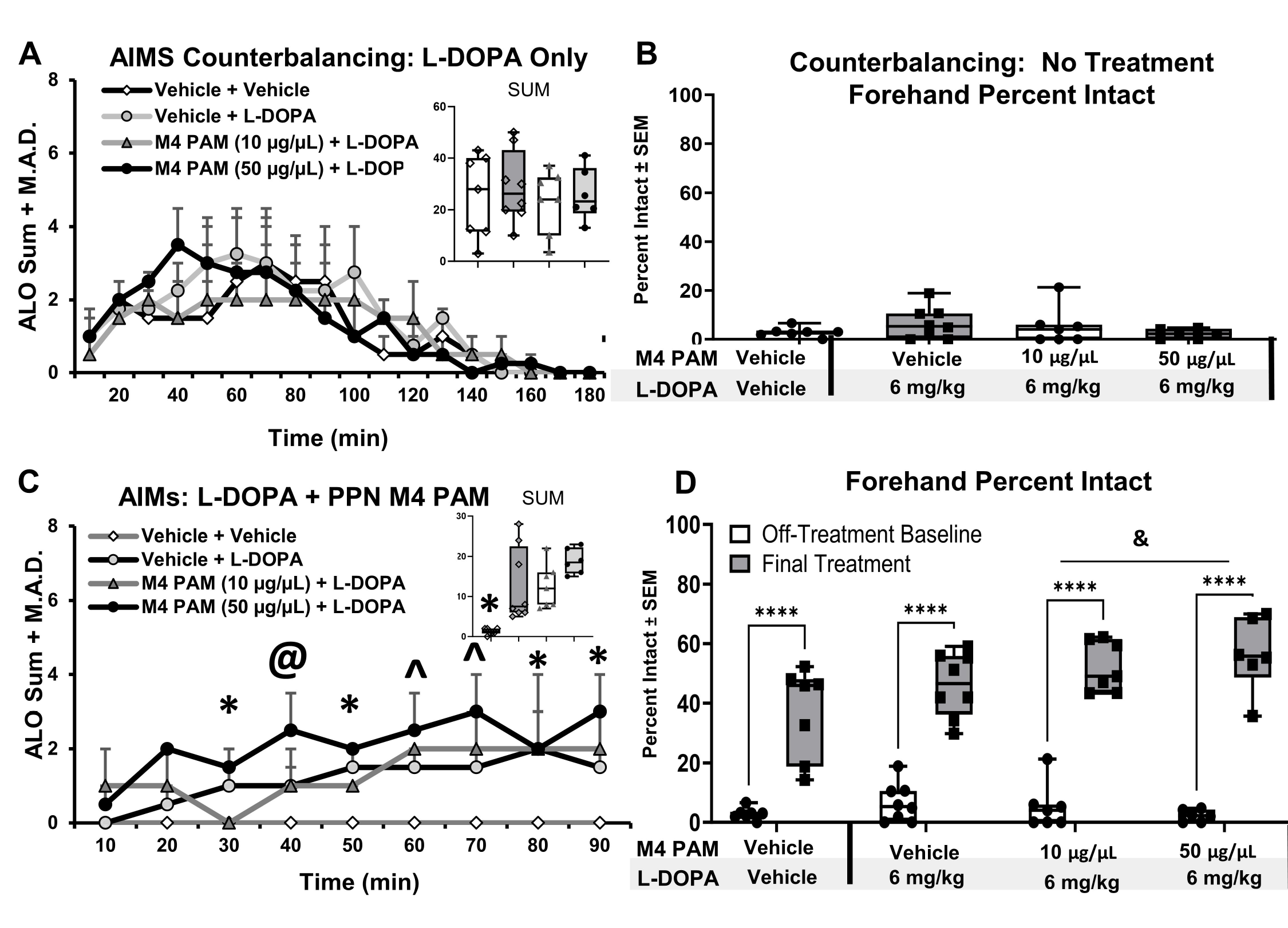
Effect of M_4_ PAM on L-DOPA-Induced Dyskinesia and Forepaw Steps. (a) L-DOPA-induced dyskinesia was scored according to the abnormal involuntary movements (AIMs) scale. (a) AIMs data from days 8 and 14 of treatment that were used for counterbalancing and placing animals into final treatment groups. Scores are shown as medians of axial, limb, and orolingual (ALO) behaviors plus the median absolute deviation (M.A.D.) for all time points. The inset graph represents the median of the overall sum of A.L.O. behaviors for each treatment group. Data were analyzed using Kruskal-Wallis non-parametric tests and showed no significant differences between final treatment groups. (b) Forepaw percent intact scores (represented in the graph as percent intact + S.E.M) were also used for counterbalancing final treatment groups. Forepaw percent intact values are derived by taking the number of steps from the right forepaw divided by the left forepaw and multiplying that value by 100. Data were analyzed using a 1-way ANOVA. There are no significant differences in baseline motor deficit. (c) Summary of all Axial, Limb, and Orolingual (A.L.O.) behaviors following pedunculopontine nucleus-(PPN) targeted microinfusion of VU0467154 (vehicle, 10 µg/µL, or 50 µg/µL) + L-DOPA (vehicle or 6 mg/kg). Data are displayed as medians of A.L.O. behaviors for each time point for each group. The inset graph represents the median of the overall sum of A.L.O. behaviors for all time points for each treatment group. Kruskal Wallis tests with Dunn post-hocs revealed that every group displayed more severe dyskinesia than the Vehicle + Vehicle group, but there were no other differences in overall sum of dyskinetic behaviors (ALO sum). There were differences at specific timepoints between 30 through 90 min. including * : p < 0.05 Vehicle + Vehicle vs. all, @ p < 0.05 Vehicle + Vehicle vs. to 50 µg/µL + L-DOPA, ∧ p < 0.05 Vehicle + Vehicle vs 10 µg/µL + L-DOPA. (d) Summary of forepaw percent intact scores on the final day of treatment + post-surgical baseline (represented in this graph as collapsed across all treatments because statistical analysis showed no significant differences). One-way ANOVA revealed that both 10 and 50 µg/µL showed a significant increase in stepping from Vehicle + Vehicle. **** p < 0.05 off-treatment baseline vs. ALL; & p < 0.05 50 µg/µL + L-DOPA and 10 µg/µL + L-DOPA vs. Vehicle + Vehicle.

**Figure 3.**
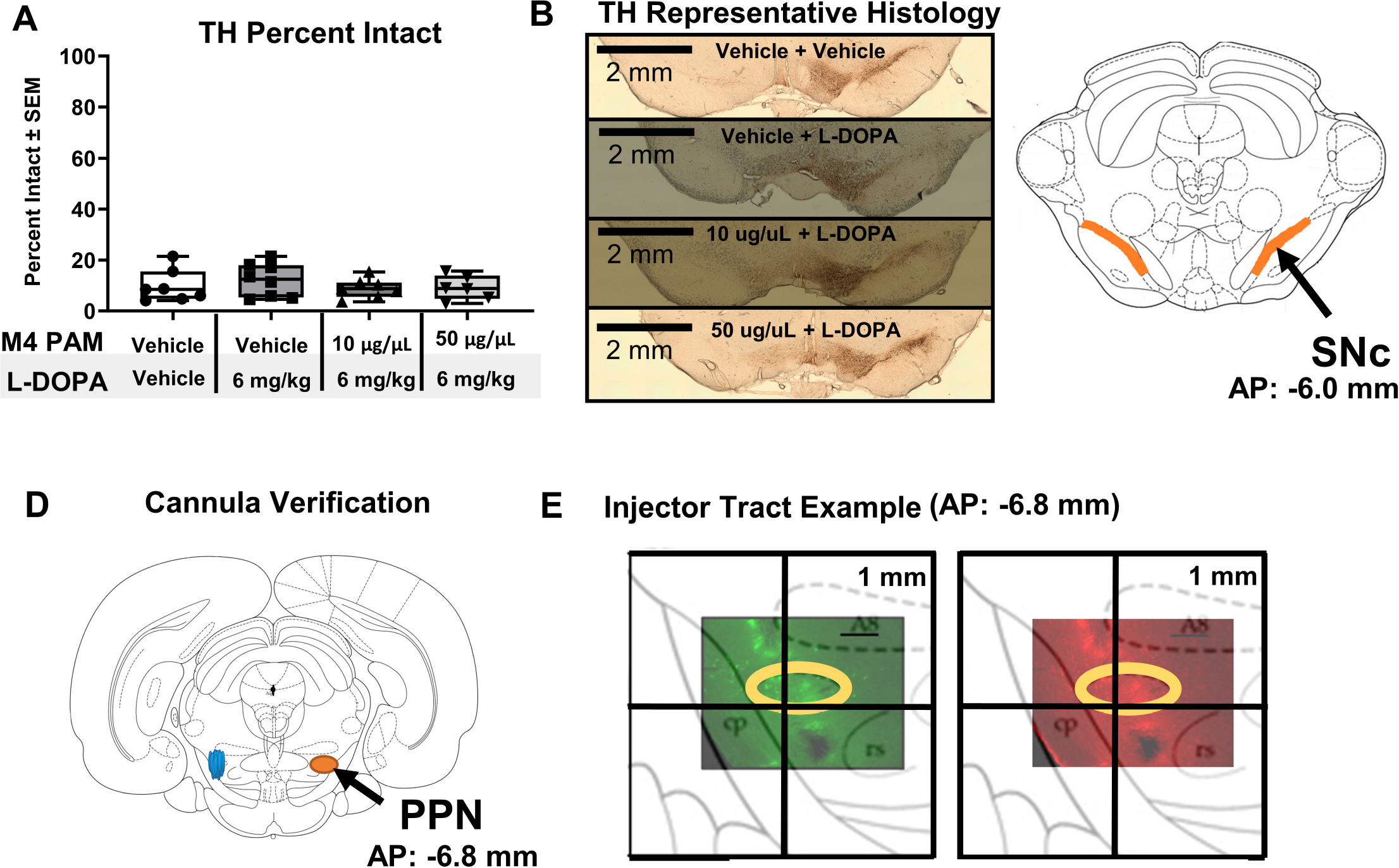
Tyrosine Hydroxylase Staining and Cannula Placement Verification. (a) Tyrosine hydroxylase (TH) Percent Intact is shown for all groups, which is calculated as total number of substantia nigra pars compacta (SNc) dopamine neurons on the left side, divided by the left side, multiplied by 100. Data were analyzed via 1-way ANOVA. All final treatment groups showed around 10% remaining dopamine neurons, and did not differ statistically. (b) TH representative histology, showing dopamine neurons of the left (lesioned) and right (intact) side for each treatment group. (c) An atlas image (Paxinos & Watson) showing the SNc in the region where dopamine neurons were counted. (d) A representative atlas image showing cannula placement in all of the animals used in this study. All areas of injection are shown as blue ovals, all animals included in this study had injections within the boundaries of the PPN. The PPN is shown on the right as an orange circle. (e) A representative image showing what the injection area looks like, with the PPN circled in the yellow oval. A 2 x 2 mm square from the relevant atlas page is overlaid over the image to show orientation of the injection. Overall, these are examples of sections which could not be used for immunohistochemical staining due to the fluorescence of the fluid in the injection site which can be seen in both green and red. Tissue damage from the injection was generally limited to about half of the PPN sections in the rostral PPN. Above the PPN that there is a tract from the injector coming down into the brain and some fluid that leaked out around the injector tract. One animal was excluded from the 50 μg/ μL group due to excessive damage from the injector that spanned the entirety of the rostral PPN.

**Figure 4.**
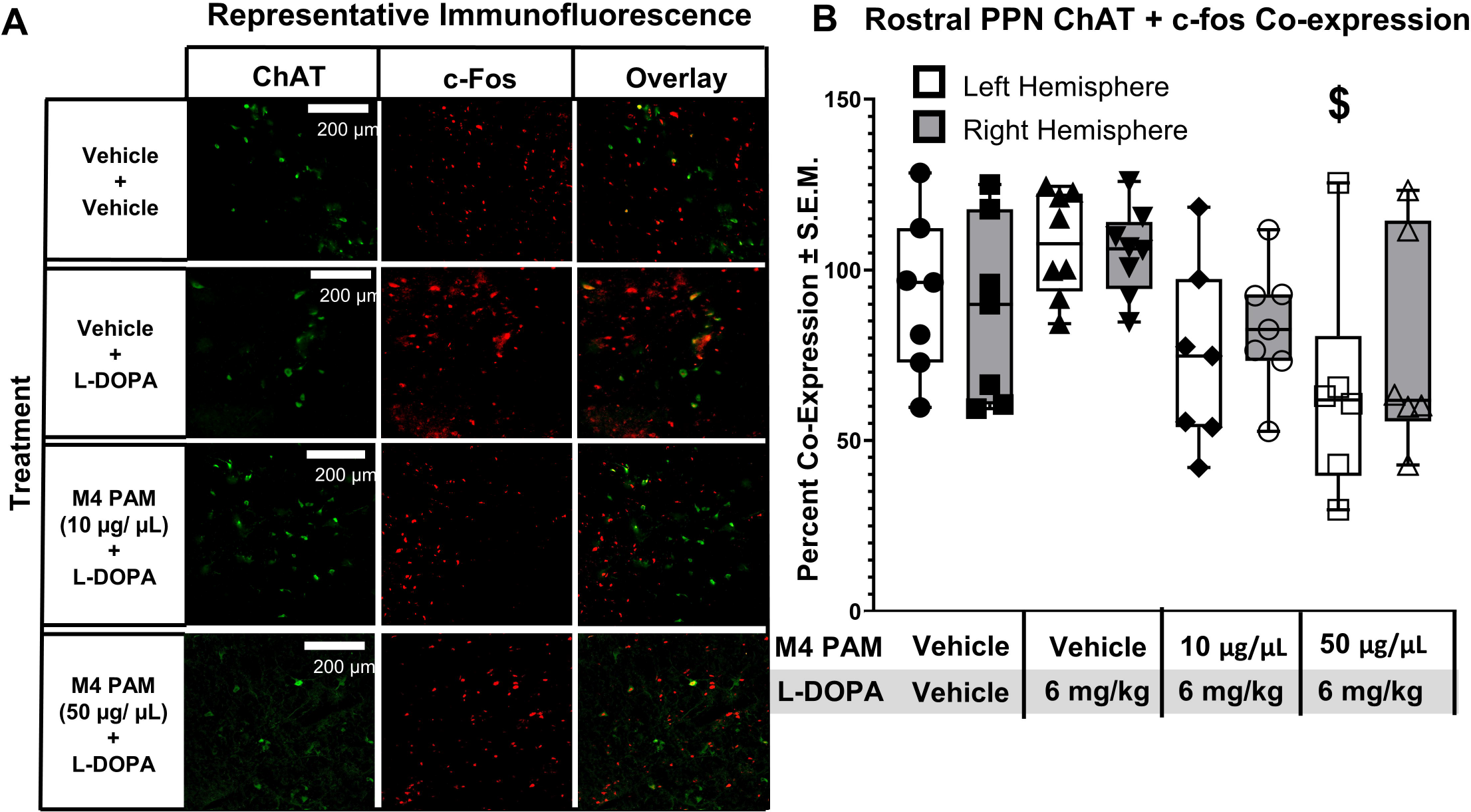
Choline Acetyltransferase and c-fos Co-expression. (a) Representative images of immunofluorescent staining on the left (lesioned) hemisphere for each group with choline acetyltransferase (ChAT) in green, c-fos in red, and the overlay in which yellow neurons represent those with co-expression. (b) Rostral PPN ChAT co-expression is quantified, showing that overall the groups receiving M_4_ PAM show less co-expression than other groups. Data are represented as the percent co-expression (calculated as ([(Total number of ChAT positive cells / c-fos positive neurons) + (co-localized neurons / total number of ChAT positive cells)] * 100). Data were analyzed using a mixed-model repeated-measures ANOVA with within-subjects factor of hemisphere and between-subjects factor of final treatment with Tukey post-hocs conducted as appropriate. $ p < 0.05 Vehicle + L-DOPA vs. 50 µg/µL + L-DOPA.

**Table 1.**
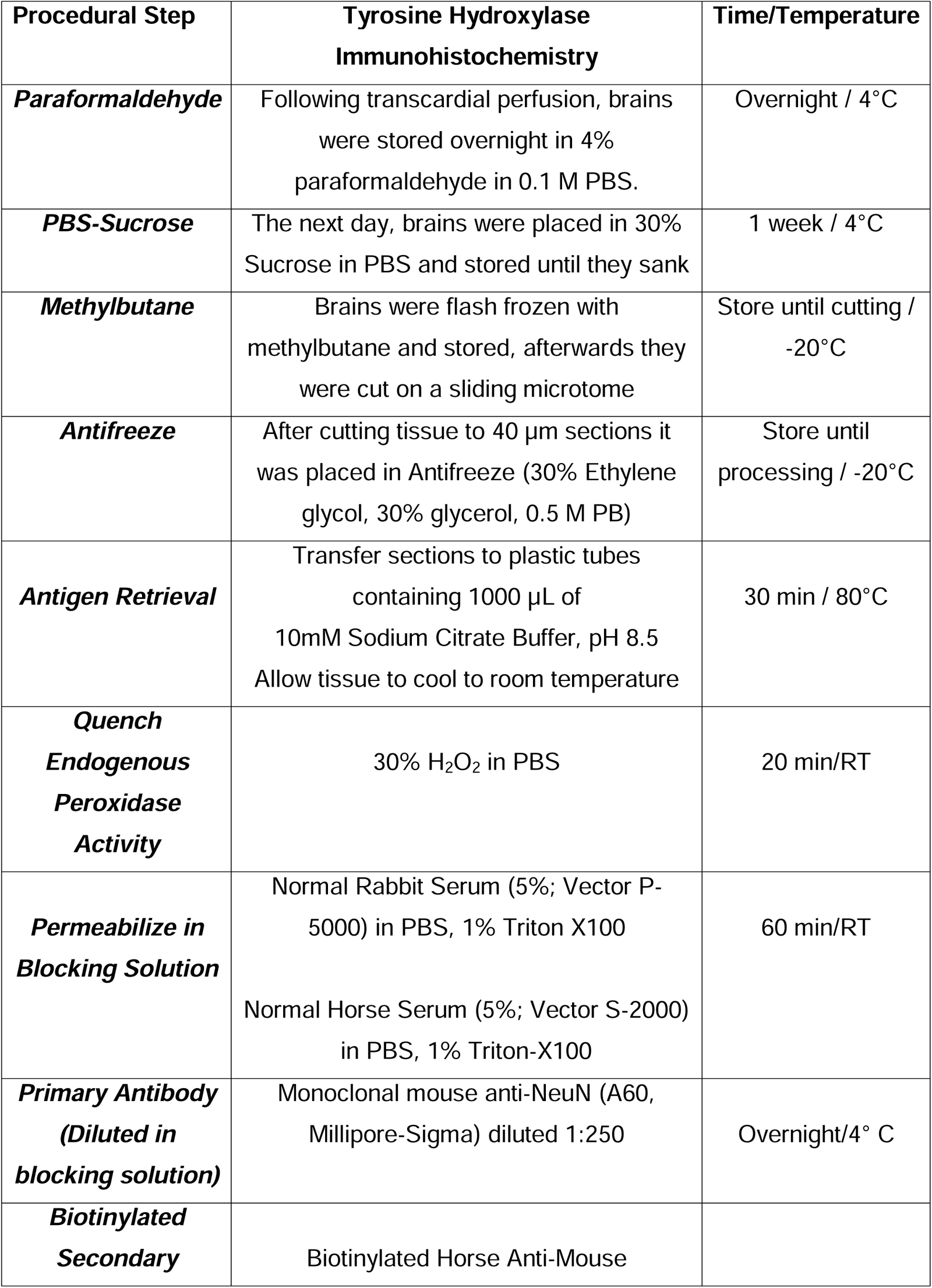
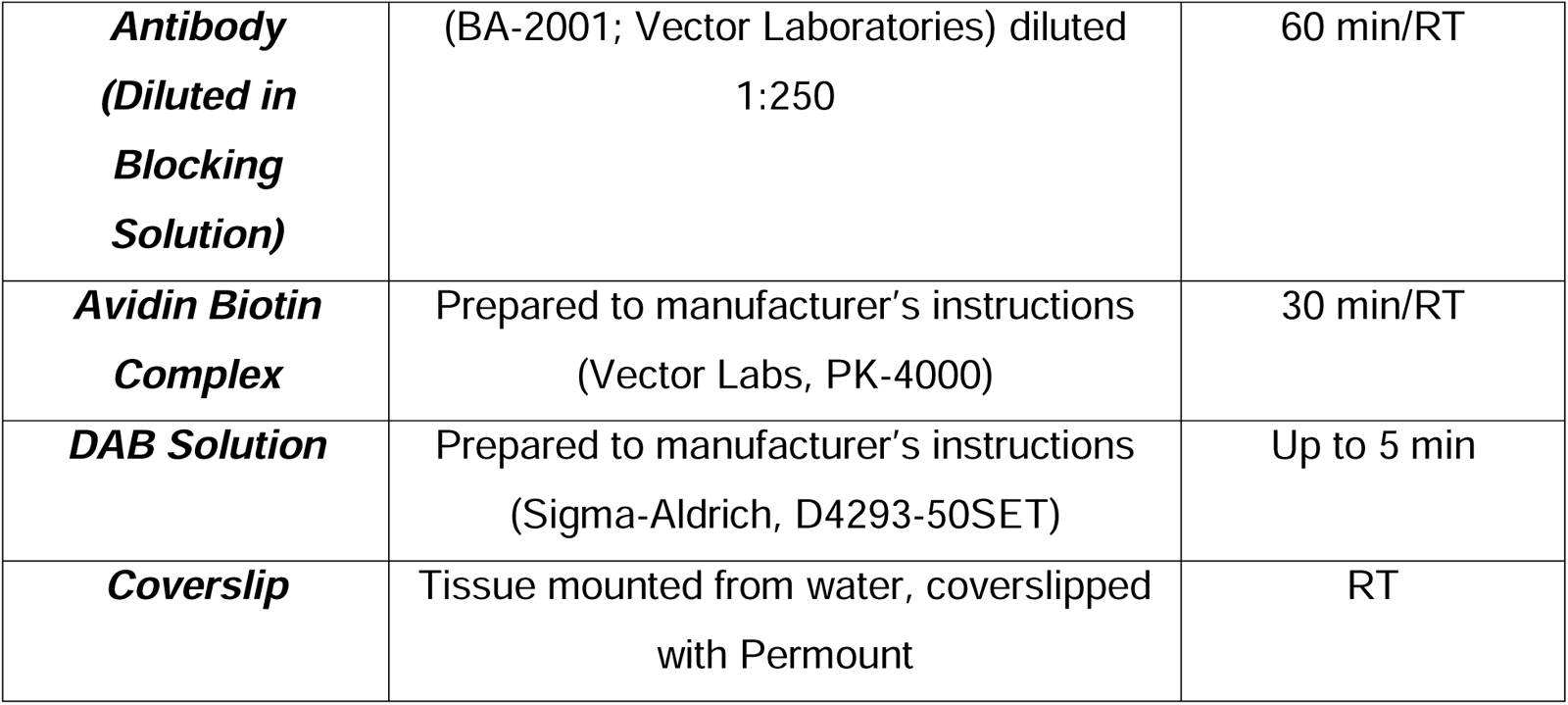
Tyrosine Hydroxylaxse Immunohistochemistry (IHC) Procedures and Reagents. This table shows a step-by-step guide to our IHC procedures with information on the reagents that were employed for tyrosine hydroxylase (TH). Please note that 0.01 M phosphate buffered saline (PBS) was used throughout unless otherwise specified. Additionally, three 5 min PBS washes were used before the first step and between each subsequent step. After the antigen retrieval, tissue was placed into net wells in a 12 well plate and processed through subsequent steps. PB: phosphate buffer; RT: room temperature (22-23°C)

**Table 2.** Choline Acetyltransferase and c-fos Immunofluorescent Immunohistochemistry Procedures and Reagents. This table shows a step-by-step guide to our Immunofluorescent (IF) staining procedures with information on the reagents that were employed for each protein of interest, including choline acetyltransferase (ChAT) and c-fos. Please note that 0.01 M phosphate buffered saline (PBS) was used throughout unless otherwise specified. Additionally, three 5 min PBS washes occurred before the first step and between each subsequent step. After antigen retrieval, tissue was placed into net wells in a 12 well plate and processed through subsequent steps. PB: Phosphate buffer; RT: room temperature (22-23°C); TBS: Tris Buffered Saline (0.01 M)

| **Procedural Step** | **ChAT + c-fos Dual Immunofluorescence** | **Storage** |
| --- | --- | --- |
| Paraformaldehyde | Following transcardial perfusion, brains were stored overnight in 4% paraformaldehyde in 0.1 M PBS. | Overnight/ 4°C |
| PBS-Sucrose | The next day, brains were placed in 30% sucrose in PBS and stored until cutting on a sliding microtome microtome | 1 month/4°C |
| Antifreeze | After cutting tissue in 40 µm sections, it was placed in Antifreeze (30% Ethylene glycol, 30% glycerol, 0.5 M PB) | Store until processing/-20°C |
| Antigen Retrieval | Transfer sections to plastic tubes containing 1000 µl of 10 mM Sodium Citrate Buffer, pH 8.5<br>Allow tissue to cool to room temperature | 30 min/80°C |
| Glycine | Incubate sections in 0.15 M glycine dissolved in TBS | 2 hours/RT |
| Permeabilize in Blocking Solution | 10% Normal Donkey Serum (Jackson ImmunoResearch 017-000-121) and 0.1% Triton-X-100 in PBS | 30 min/RT |
| Primary Antibody Cocktail | Incubate sections in primary antibody cocktail in blocking solution consisting of monoclonal rabbit anti-c-Fox antibody (1:100 dilution; Cell Signaling, 2250) and goat anti-ChAT antibody (1:100 dilution, Millipore-Sigma) | Overnight/4°C |
| Secondary Antibody Cocktail | Incubate sections in a secondary antibody cocktail in blocking solution consisting of AlexaFluor 488 AffiniPure Donkey Anti-Goat (1:1000 dilution; Jackson ImmunoResearch, 705-545-003) and Alexa Fluor 594 AffiniPure Donkey Anti-Rabbit (1:1000 dilution, Jackson ImmunoResearch 715-585-152). | Overnight/4°C |
| Mount Slide with Gelvatol Mounting Medium | Tissue was mounted onto slides with Gelvatol Mounting Medium and left to dry in a drawer overnight before being placed in a light-protected slide box. | 5 min/RT |

### 2.3 Surgery

Rats were anesthetized and stabilized in a Kopf stereotax (as described in Chambers et al., 2019). A craniotomy was performed above the lesion site and then 6-OHDA (12 µg in 4 µL) was infused into the left medial forebrain bundle (From bregma ML: -2.0 mm; AP: -1.8 mm; DV: -8.6 mm). During the same surgery, rats received a unilateral PPN-targeted cannula (from bregma AP: -6.8 mm; ML: -2.0 mm; from dura DV: -4.0 mm). Buprinex (0.03 mg/kg, i.p.) was administered prior to surgery and every 12 h for 24 h following surgery. Rats received one wound clip behind the cannula to promote tissue recovery. Fifty-four rats received surgery for this study, but 14 of these rats died or were euthanized following surgery due to postoperative complications related to cannula placement above the brainstem such as blood clots, swelling, and failure to thrive after surgery.

### 2.4 Forepaw Adjusting Steps (FAS)

Forepaw stepping deficits and L-DOPA motor improvement were measured using the FAS test. Rats were dragged laterally across a surface at a rate of 90 cm over 10 s while the experimenter restrained all paws except for one forepaw as previously described (Chambers et al., 2019; Chang et al., 1999). Adjusting steps made by the forepaw were counted by a trained rater blind to condition. Three trials were performed for each forepaw for both backward and forward stepping. For our analysis we used just the foreward stepping values for each paw. A forepaw percent intact score was calculated as follows: total right forepaw steps / total left forepaw steps * 100. Two animals had over 50% intact stepping, which shows an incomplete lesion and thus were excluded from the study.

### 2.5 Abnormal Involuntary Movements Scale (AIMs)

Dyskinetic behaviors were quantified using the AIMs test (Cenci & Lundblad, 2007; Putterman et al., 2007). Rats were administered L-DOPA methyl ester (6 mg/kg, s.c.) formulated with 15 mg/kg of the peripheral decarboxylase inhibitor Benserazide dissolved in 0.9% saline and 0.1% ascorbate (hereafter L-DOPA). Thereafter, rats were placed in plexiglass cylinders and dyskinesia was rated for 1 min every 10 min for 90 min beginning 10 min after L-DOPA administration. Axial, limb, and orolingual (ALO) behaviors were scored on a scale of 0 to 4 as previously described (Chambers et al., 2019; 2021). All remaining animals displayed dyskinesia and were retained for microinjection (n = 38).

### 2.6 PPN Microinjection

Prior to testing rats were habituated to restraint and the microinjection apparatus. An 18.8 mm injector connected to a 10 µL Hamilton syringe (Hamilton Company, Reno, NV) by polyethylene tubing (0.46 mm in diameter; PlasticsOne) was inserted slowing into the guide cannula. Rats received a 2 min infusion (0.25 µL/min; 0.5 µL total volume). Rats received either vehicle, 10, or 50 µg/µL of VU0467154. The injector remained in place for 3 min to allow for absorption, after which it was slowly removed, the obdurator replaced, and the rat was then injected with L-DOPA (6 mg/kg, s.c.). Rats received only one microinfusion before being euthanized 90 min after L-DOPA administration.

### 2.7 Double-Labeling Immunofluorescence for ChAT and c-fos

To quantify colocalization of ChAT and c-fos, at least 2 coronal sections containing rPPN were used for immunofluorescence (From bregma: AP -6.72 to AP -7.3 mm; see Table 1 for Procedure and reagents). To allow for accurate quantification we chose slices that did not show damage from the injector or autofluorescence from the injected drug (See figure 3D-3E). Images were acquired on a Keyence BZ-X series microscope at 20x magnification using 2 separate channels (GFP for ChAT and Texas Red for c-fos). Each channel was z-stacked at 0.3 pitch and focused prior to image acquisition. Images were processed and analyzed in ImageJ by an investigator blind to condition. The rollerball tool (set to 50%) was used to subtract background, brightness was uniformly adjusted, and images were sharpened to aid in analysis. Green and red images were then fused to create an overlay. Percentage of colocalization was calculated for each subject by the following equation: ([(Total number of ChAT positive cells / c-fos positive neurons) + (co-localized neurons / total number of ChAT positive cells)] * 100). These colocalization values were averaged for slices on each side for each subject before data analysis. PPN slices were also used to verify successful cannula placement. Of the 38 animals receiving microinjections, 3 were excluded due to injection not falling within the boundaries of the PPN, 1 animal was excluded because of excessive tissue damage from the PPN injection site (over half of the rostral PPN damaged), and 4 animals were excluded due to a drug dosing error.

### 2.8 TH Immunohistochemistry

To verify 6-OHDA lesion, (as described in Chambers et al., 2021) TH immunohistochemistry was employed on at least 2 coronal substantia nigra pars compacta (SNc) sections per animal (From bregma: AP -5.2 to AP -5.8 mm; See Table 1 for Procedure and Reagents). Nigral brain slices with TH staining were imaged with a BZ-X series microscope at 10x magnification. For TH cell counts total enumeration was used (Chambers et al., 2021; Sellnow et al., 2019). Data are represented as percent intact values, calculated as: [(left (lesioned) cell count / right (intact) cell count) * 100]. Of the 30 animals receiving microinjections which had successful placement of the injection within the PPN, 2 were excluded due to TH counts above 40%, indicating incomplete lesion, leaving 28 animals remaining for data analysis.

### 2.9 Data Analysis

For all experiments, data were analyzed using GraphPad Prism version 10.0.03; α = 0.05. ALO AIMs data were analyzed for both: 1) the overall sum of axial, limb, and orolingual (ALO) behavior; and 2) ALO sums for each given time point. AIMs data were analyzed using Kruskal-Wallis non-parametric testing with Dunn’s post-hoc tests as needed to analyze effects of treatment. FAS data for counterbalancing were analyzed with a one-way ANOVA with final treatment group as a factor. Data from the final test day were first analyzed via a 2-way mixed-model repeated-measures ANOVA with within-subjects factor of test day (2: post-surgical baseline (off treatment), microinfusion test day), and then for between-subjects factor of final treatment (4: Vehicle + Vehicle, Vehicle + L-DOPA, 10 µg/µL VU0467154 + L-DOPA, 50 µg/µL VU0467154 + L-DOPA).

Percent colocalization of ChAT and c-fos were analyzed via a mixed-model ANOVA with a within-subjects factor of hemisphere (2: lesioned, unlesioned), and a between-subjects factor of treatment (4). Percent intact values for TH IHC were measured via 1-way ANOVA with between-subjects factor of treatment (4). Tukey post-hocs were employed to correct for multiple comparisons as appropriate for all ANOVAs. T-tests measuring the difference between TH immunoreactivity in the left (lesioned) vs. right (intact) hemisphere were also performed.

## 3.0 Results

### 3.1 AIMs

After 2 weeks of daily L-DOPA treatment, median ALO sums from days 8 and 14 were calculated, and rats were divided into equally counterbalanced treatment groups with no difference in overall ALO sums or timepoint differences in ALO scores (Figure 2A). Following two days of drug washout, rats received 1 PPN-targeted microinfusion of VU0467154 (vehicle, 10, or 50 µg/µL) 5 min before L-DOPA (vehicle or 6 mg/kg). Analysis of ALO sums showed an effect of treatment on overall dyskinesia severity, H(3) = 17.82, p < 0.05. Dunn’s post hocs revealed that all groups treated with L-DOPA showed more severe dyskinesia than the Vehicle + Vehicle group p < 0.05. Time course effects were also present at the 20 through 90 min time points. Dunn’s post hocs showed that Vehicle + Vehicle group had more dyskinesia than the 50 µg/µL + PAM group for all timepoints in this range. Furthermore, at the 50 and 80 min time point Vehicle + L-DOPA displayed more dyskinetic behaviors than the Vehicle + Vehicle group. Finally, at the 90 min time point, 10 µg/µL + L-DOPA showed more dyskinesia than the Vehicle + Vehicle group. Neither of the PAM-treated groups were different than the Vehicle + L-DOPA group in the overall analysis of ALO sums or in the time course analysis, suggesting that M_4_ in the rostral PPN do not affect LID.

### 3.2 FAS

Two weeks after surgery, FAS was performed off-treatment to determine baseline motor impairment. At baseline, a 1-way ANOVA analysis revealed no statistical difference between groups, indicating that treatment groups had equal levels of motor impairment (Figure 2B). A 2-way mixed-model repeated-measures ANOVA was used to examine within subjects factors of test day (2: off treatment baseline, final test day), and between-subjects factor of final treatment (4: Vehicle + Vehicle, Vehicle + L-DOPA, 10 µg/µL + L-DOPA, or 50 µg/µL + L-DOPA). Although there was no interaction between final treatment and test day, there was a significant main effect of test day, F(1,25) = 300.20, p < 0.05, ηp22 = 0.92. Overall, rats had increased forepaw stepping on the lesioned side on the final test day than they did at the off-treatment baseline. There was also a main effect of final treatment, F(3,25) = 3.47, p < 0.05, ηp22 = 0.96. Overall, animals receiving 10 or 50 µg/µL had significantly increased stepping as compared to the Vehicle + Vehicle group (Figure 2D), whereas animals treated with Vehicle + L-DOPA did not show significantly increased stepping compared to this group, suggesting a small additional motor benefit of M_4_ PAM microinfusion with L-DOPA above the motor benefit of L-DOPA alone.

### 3.3 ChAT + c-fos Co-Expression

We conducted dual ChAT and c-fos IHC in the rPPN to understand how cholinergic cell activity corresponds to treatment. A repeated-measures ANOVA with between-subjects factor of hemisphere (2: left, right) and between-subjects factor of treatment (4: Vehicle + Vehicle, Vehicle + L-DOPA, 10 µg/µL + L-DOPA, 50 µg/µL + L-DOPA) showed no effect of hemisphere on c-fos expression, but a significant effect of treatment, F(3,25) = 3.26, p < 0.05. Tukey post-hocs revealed effects of treatment, F(3, 25) = 4.79, p < 0.05, ηp2 = 0.36. Specifically, on the left (lesioned) side the Vehicle + L-DOPA group had a higher percentage of ChAT neurons coexpressing c-fos as compared to group injected with 50 µg/µL + L-DOPA, p < 0.05, suggesting that M_4_ PAM infusion may normalize PPN activity during LID.

### 3.4 TH Immunoreactivity

In order to measure lesion efficacy we counted TH+ cells in the substantia nigra in at least 2 coronal slices per animal. Data were first analyzed as percent intact values via 1-way ANOVA. Overall, there was no significant difference between TH percent intact in the treatment groups, (M_Vehicle + Vehicle_ = 9.88, SEM_Vehicle + Vehicle_ = 2.41; M_Vehicle + L-DOPA_=14.30, SEM_Vehicle + L-DOPA_ = 2.95; M_10 µg/µL + L-DOPA_ = 8.90; SEM10 _µg/µL + L-DOPA_ = 1.45; M_50µg/µL + L-DOPA_ =9.17, SEM_50 µg/µL L-DOPA_ = 1.93), indicating statistically similar lesions for each group. We then collapsed across conditions and performed a paired-sample t-test on left (lesioned) and right (intact) hemisphere, which revealed, as expected, that there was a significant difference in TH immunoreactivity between intact and lesioned hemispheres, t(28)=16.51, p < 0.05, demonstrating successful 6-OHDA lesion.

## 4.0 Discussion

PPN contribution to L-DOPA-mediated behaviors remains under-investigated in PD. Therefore, we used the hemiparkinsonian rat model of PD to investigate the effects of PPN infusion of M_4_ PAM VU0467154 on LID, L-DOPA-mediated motor benefit, and c-fos expression in PPN cholinergic neurons. We hypothesized that PPN infusion of M_4_ PAM would decrease LID severity, forepaw stepping, and c-fos expression in PPN cholinergic neurons. Similar to prior research (Chambers et al., 2021), our IHC analysis demonstrated successful lesion, with an average of 90% reduction in SNc DA neurons. Overall, M_4_ PAM augmented L-DOPA’s effects, increasing forepaw stepping on L-DOPA and non-significantly increasing dyskinesia. Additionally, c-fos expression in PPN cholinergic neurons was increased in rats injected with L-DOPA, and was reduced by M_4_ PAM. This suggests that PPN cholinergic neurons could be a unique therapeutic target for augmenting L-DOPA-derived motor benefit without affecting LID. Additionally, when combined with our previous findings (Chambers et al., 2021) that PPN cholinergic neurons influence freezing of gait and aspects of gait related to balance and stability, it is possible that PPN cholinergic neurons may also be able to modify PD gait deficits without negatively impacting LID or L-DOPA-mediated motor benefit.

Stimulation of PPN cholinergic neurons improves forepaw akinesia in a PD model by normalizing nigrostriatal DA transmission (Pienaar et al., 2015; Sharma et al., 2020). Excitation of PPN cholinergic terminals in the SNc also increases locomotor behavior in normal animals (Xiao et al., 2016). Given the afore-mentioned pro-motor effects of stimulating PPN cholinergic neurons (however, see also Takakusaki et al., 2003; Sherman et al., 2015; and Roseberry et al., 2016), we expected that inhibition with M_4_ PAM would reduce L-DOPA-mediated motor improvement. Instead, we observed that M_4_ PAM increased forepaw stepping on L-DOPA. Although our Vehicle + L-DOPA group was not significantly different than Vehicle + Vehicle, our 10 µg/µL + L-DOPA and 50 µg/µL + L-DOPA groups did show a significantly higher level of stepping than Vehicle + Vehicle, suggesting that M4 PAM may slightly augment L-DOPA’s motor effects. Critically, our manipulation is not selective for specific PPN neurons or projections. Instead, our results show the net effect of M_4_ signaling in the rPPN which contains mainly cholinergic and GABAergic neurons. Although both types of neurons may contain M_4_, PPN cholinergic neurons have more dense projections to other brain areas, suggesting that our effect is likely mediated by cholinergic efferents, particularly those traveling to the striatum, substantia nigra pars reticulata (SNr) and SNc (Chambers et al., 2020). Within the striatum, ACh release has been shown to oppose effects of DA through M_4_ located on medium spiny neuron cell bodies (Shen et al., 2015). Similarly, ACh release from PPN terminals in the SNr opposes the effects of DA by inhibiting terminals of D1-bearing MSNs through M_4_ (Moehle et al., 2017). Within the SNc, PPN cholinergic terminals release ACh which binds to nAChR (Liu et al., 2022) and M5 (Foster et al., 2014), eliciting an excitatory effect on DA neurons. Prior studies demonstrating modulation of PD motor deficits by PPN cholinergic projections to the SNc have used the lactacystin model of PD which is associated with less severe DA loss. In our severe 6-OHDA lesion model, it is questionable whether enough SNc DA neurons remain to observe this effect. Beyond cholinergic influences on movement, rigorous work from other groups suggest that PPN and neighboring cuneiform nucleus glutamatergic neurons play a larger role in locomotor behavior (Takakusaki et al., 2003; Sherman et al., 2015; Roseberry et al., 2016); however, the role of glutamatergic PPN neurons in PD models remains underinvestigated and is beyond the scope of this study. Finally, the fact that the group receiving Vehicle + Vehicle did not show differences in forepaw stepping with respect to the Vehicle + L-DOPA group requires further comment. Both groups received two weeks of chronic treatment with L-DOPA so that we could create equally lesioned, equally dyskinetic final treatment groups. Even after two days of drug washout, forepaw stepping remained elevated in the Vehicle + Vehicle group. This is likely explained by long-term changes in the basal ganglia circuit which are caused by L-DOPA and which have been observed by others in the clinc (Albin & Leventhal, 2017; Cilia et al., 2020; Falkenburger et al., 2022) and in preclinical models (Kang & Auinger, 2012). This may mean that modulating PPN cholinergic activity may be efficacious as a strategy to augment L-DOPA effects in late-stage PD. If the Vehicle + L-DOPA group had been L-DOPA naïve, it is likely that we would have observed larger differences between groups both for forehand stepping and for c-fos expression. Therefore, we also compared final treatment day with the initial off-treatment baseline to better recapitulate forepaw stepping behavior in L-DOPA naïve animals (Figure 2D).

We also examined modulation of LID as prior research shows that M_4_ PAM can lessen LID in preclinical models (Shen et al., 2015) and that the PPN may contribute to motor stereotypy. Our prior work (Chambers et al., 2019) suggests that ACh receptor antagonism in the PPN slightly increases LID and forepaw stepping on L-DOPA. Additionally PPN infusion of general muscarinic antagonists causes motor stereotypy in normal rats, which is reversed by general muscarinic agonists (Mathur et al., 1997). Given that systemic injection of M_4_ PAM reduces dyskinesia and that muscarinic antagonism of the PPN worsens dyskinesia and motor stereotypy, we expected to find that M_4_ PAM infusion lessened LID. On the contrary, we found that M_4_ PAM non-significantly increased LID (Figure 2C). The slight increase that we observed in ALO behaviors may correspond to a decrease in ACh release in either the striatum or SNr, thus disinhibiting direct pathway neurons or terminals, respectively.

Multiple studies have shown an increase in PPN IEG expression in preclinical PD models chronically receiving L-DOPA (Bensaid et al., 2016; Ding et al., 2007), suggesting that PPN neuron activity is increased by L-DOPA treatment. Currently no studies have investigated which specific subtypes of PPN neurons may be activated by L-DOPA. Given the pro-motor effects of stimulating rPPN cholinergic neurons (Dautan et al., 2014; 2016; Xiao et al., 2015; however see also Moehle et al., 2017), we investigated whether these specific neurons show an increase in IEG expression in response to L-DOPA. In accordance with our hypothesis, we found that the expression of the IEG c-fos is slightly, but non-significantly increased in PPN cholinergic neurons after systemic L-DOPA administration. Furthermore, treatment with M_4_ PAM decreased c-fos expression in PPN cholinergic neurons, and significantly reduced c-fos expression on the left side relative to the rats treated with Vehicle + L-DOPA. This suggests that it is PPN cholinergic neurons specifically which are activated by L-DOPA treatment. The fact that there was no significant difference between Vehicle + Vehicle and Vehicle + L-DOPA groups also requires further comment. Like the forehand stepping on the final test day, the lack of observed difference between the Vehicle + Vehicle and the Vehicle + L-DOPA group may be because Vehicle + Vehicle animals were previously treated with L-DOPA prior to the final treatment day. If we had included an L-DOPA naïve group, we likely would have observed a significant elevation in c-fos expression with Vehicle + L-DOPA. Future studies should be conducted to more closely examine changes in PPN cholinergic activity in response to L-DOPA administration. However, despite this limitation, our results suggest that PPN cholinergic neurons upregulate their activity in response to L-DOPA and that M_4_ PAM normalizes cholinergic activity in response to L-DOPA.

This investigation adds to a growing body of literature suggesting a role for PPN cholinergic neurons in improving motor deficits in preclinical models and in increasing the motor benefit of L-DOPA. Taken together, our results suggest that PPN cholinergic neurons may be a promising therapeutic target for dose-sparing strategies with L-DOPA because although they modify parkinsonian motor deficits, they do not significantly affect LID in this study or in our other studies (Chambers et al., 2019; 2021). Future studies should dissect the effects of specific PPN cholinergic projections on PD motor deficits and L-DOPA-mediated behaviors, with the goal of identifying a projections which contribute maximally to these effects. Overall, it is likely that targeting just one specific PPN cholinergic projection would offer a promising therapeutic strategy for augmenting the effects of L-DOPA, improving both PD motor and gait deficits without worsening LID.

## List of Abbreviations

6-OHDA: 6-hydroxydopamine
ACh: Acetylcholine
AIMs: Abnormal involuntary movements scale
ALO: Axial, limb, and orolingual
ChAT: Choline acetyltransferase
DA: Dopamine
D1: Dopamine D1 receptor
FAS: Forepaw adjusting steps
GABA: Gamma-aminobutyric acid
IHC: Immunohistochemistry
IR: Immunoreactivity
LID: L-DOPA-induced dyskinesia
M4: M4 muscarinic acetylcholine receptor
PBS: Phosphate buffered saline
PD: Parkinson’s disease
PPN: Pedunculopontine nucleus
rPPN: Rostral pedunculopontine nucleus
TH: Tyrosine hydroxylase

## Author Contributions

Participated in research design: Chambers, Bishop

Conducted experiments: Chambers, Mclune, Coyle, Sergio, Delpriore, Lanza Performed data analysis: Chambers, Bishiop

Wrote or contributed to writing of manuscript: Chambers, Bishop, Resources (Made and provided VU0467154): Lindsley, Conn

## Data Availability

Data from this manuscript will be available on the Mendeley data repository following peer-reviewed publication

